# Guest-host supramolecular assembly of injectable hydrogel fibers for cell encapsulation

**DOI:** 10.1101/2021.02.26.430926

**Authors:** Beverly Miller, Audrey Hansrisuk, Christopher B. Highley, Steven R. Caliari

## Abstract

The fibrous architecture of the extracellular matrix (ECM) is recognized as an integral regulator of cell function. However, there is an unmet need to develop mechanically robust biomaterials mimicking nanofibrous tissue topography that are also injectable to enable minimally invasive delivery. In this study we have developed a fibrous hydrogel composed of supramolecularly-assembled hyaluronic acid (HA) nanofibers that exhibits mechanical integrity, shear-thinning, rapid self-healing, and cytocompatibility. HA was modified with methacrylates to permit fiber photocrosslinking following electrospinning and either ‘guest’ adamantane or ‘host’ β-cyclodextrin groups to guide supramolecular fibrous hydrogel assembly. Analysis of fibrous hydrogel rheological properties showed that the mixed guest-host fibrous hydrogel was more mechanically robust (6.6 ± 2.0 kPa, storage modulus (G′)) than unmixed guest hydrogel fibers (1.0 ± 0.1 kPa, G′) or host hydrogel fibers (1.1 ± 0.1 kPa, G′) separately. The reversible nature of the guest-host supramolecular interactions also allowed for shear-thinning and self-healing behavior as demonstrated by cyclic deformation testing. Human mesenchymal stromal cells (hMSCs) encapsulated in fibrous hydrogels demonstrated satisfactory viability following injection and after seven days of culture (> 85%). Encapsulated hMSCs were more spread and elongated when cultured in viscoelastic guest-host hydrogels compared to non-fibrous elastic controls, with hMSCs also showing significantly decreased circularity in fibrous guest-host hydrogels compared to non-fibrous guest-host hydrogels. Together, these data highlight the potential of this injectable fibrous hydrogel platform for cell and tissue engineering applications requiring minimally invasive delivery.

## Introduction

The extracellular matrix (ECM) is a complex three-dimensional (3D) microenvironment that provides mechanical support, protection, and regulatory signals to embedded cells^1,2^. Hydrogels are widely used to mimic native ECM due to their ability to exhibit tissue-like properties, including viscoelastic mechanics and high water content comparable to the tissues they are intended to model and replace^3–5^. In the context of tissue engineering, there is growing appreciation for the importance of the fibrillar architecture of ECM and its role in providing biophysical cues that regulate cell behavior^6–9^. In particular, there is special interest in robust 3D hydrogels that can mimic the fibrous networks present in many musculoskeletal tissues while allowing tuning of network biophysical and biochemical properties^10–14^. A popular method of forming mechanically robust fibrous materials is electrospinning, a simple, scalable, and cost-effective biofabrication technique that produces nanofibers with tunable physicochemical properties^15–18^. Electrospinning has also garnered interest owing to its ability to produce nanofibers, typically using synthetic polymers^19,20^, with topographical similarity to collagen and elastin fibrils within the native ECM^21–24^.

Currently, much of our understanding of how cells interact with fibrous microenvironments stems from two-dimensional (2D) culture studies of cells seeded atop electrospun fibers. Importantly, recent work has demonstrated that human mesenchymal stromal cells (hMSCs) cultured on electrospun synthetic fibers recapitulate many of the behaviors observed when cells are cultured in natural matrices like collagen, in contrast to observations of cells grown on synthetic hydrogels^25^. A separate study using a naturally-derived fibrous collagen platform showed similar results, demonstrating that hMSCs grown on more compliant collagen fibers were able to remodel their surroundings more easily, leading to increased spreading and focal adhesion organization^26^. While these studies were critical in establishing the importance of including fibrous architectural cues in biomimetic cell culture models, cell behavior in 3D microenvironments can depart dramatically from 2D culture formats^13,27^. To that end, recent findings demonstrated that the inclusion of embedded nanofibers in 3D hydrogels drives increased cell spreading and nuclear localization of the mechanoregulatory transcriptional coactivator Yes-associated protein (YAP)^28^. Further, a study utilizing multifunctional peptide-based assemblies to form mechanically robust and hierarchically-structured hydrogels found that matrices with longer fibrils facilitated hMSC morphologies similar to those observed in native collagen environments^29^. This growing body of work underscores the importance of integrating fibrous structural signals into biomaterial design.

While recreating biophysical cues, such as fibrous architecture, present in ECM has been a focus of the tissue engineering community in recent years^30^, interest in developing injectable (shear-thinning and self-healing) or *in situ* gelling materials has intensified for both clinical (including drug delivery^31^ and tissue engineering^32^ with minimally invasive delivery) and additive manufacturing (3D-printing^33^) applications. Clinically, injectable materials for tissue engineering applications offer major advantages such as reduced patient discomfort and treatment cost^34,35^. Recent advancements in synthetic chemistry and materials science have provided researchers with a library of techniques to create injectable biomaterials^32^. Injectable strategies include the use of ionically crosslinked materials^36^, jammed particles^37,38^, supramolecular polymeric materials^39^, and materials that gel *in situ* via thermoresponsive^40^ or pH-responsive^41^ self-assembly. Guest-host (e.g., adamantane-β-cyclodextrin) supramolecular chemistries are attractive for injectable hydrogel design due to their ability to support shear-thinning and self-healing behavior while also allowing careful tuning of viscoelastic mechanical properties through both primary and secondary crosslinking mechanisms^42,43^. Guest-host assembled hydrogels are also suitable cell carriers for injectable delivery since their non-Newtonian properties help protect cells from excessive shear during needle extrusion^44^. However, achieving fibrillar topographies in mechanically robust injectable hydrogels remains challenging. While self-assembling peptides can serve as injectable nanofibrous hydrogels^45^, these materials are often limited by poor long-term mechanical properties^5,8^. Electrospun hydrogel materials, as described earlier, show robust mechanics and support biomimetic cellular behaviors. However, electrospun scaffolds typically require physical implantation and are not amenable to injectable delivery^46,47^.

To address the need for a fibrous hydrogel system amenable to minimally invasive delivery, we have combined strategies in biomanufacturing, supramolecular chemistry, and tissue engineering to produce a mechanically robust injectable fibrous hydrogel capable of supporting cell encapsulation. The engineering design brings together hyaluronic acid (HA) electrospun hydrogel nanofibers, functionalized with either guest (adamantane, Ad) or host (β-cyclodextrin, CD) supramolecular moieties, that when joined together creates a shear-thinning and self-healing hydrogel fiber network via guest-host complexation. HA was utilized for its amenability to functionalization with reactive groups enabling both photocrosslinking and guest-host assembly as well as its decades long record of clinical use as a naturally-derived biopolymer^22^. In this work we assessed the architecture of the fabricated hydrogel nanofibers, their rheological properties, and their ability to support sustained cell viability following injection.

## Materials and Methods

### Materials

Sodium hyaluronate (sodium HA, 64 kDa) was purchased from Lifecore Biosciences. β-cyclodextrin (CD), hexamethylenediamine (HDA), ammonium chloride, and p-Toluenesulfonyl chloride were purchased from TCI America. Tetrabutylammonium hydroxide (TBA-OH) was purchased from Acros Organics. All other materials were purchased from Sigma-Aldrich.

### Synthesis of β-CD-HDA

Synthesis of 6-(6-aminohexyl)amino-6-deoxy-β-cyclodextrin (β-CD-HDA) was performed as previously described^34,42^, first via synthesis of the intermediate 6-o-monotosyl-6-deoxy-β-cyclodextrin (CD-Tos) using β-cyclodextrin (β-CD) and p-toluenesulfonyl chloride (TosCl). Briefly, β-CD was suspended in water and cooled to 0°C. TosCl was dissolved in minimal acetonitrile and added dropwise. After stirring at room temperature for 2 h, sodium hydroxide was added dropwise. The reaction was stirred at room temperature for 30 min before the addition of solid ammonium chloride to obtain pH 8.5. The solution was cooled on ice and the precipitate collected. The CD-Tos product was washed first with cold deionized (DI) water and then cold acetone before being dried under vacuum. After product confirmation using ^1^H NMR (500 MHz Varian Inova 500), a round bottom flask was charged with the CD-Tos, HDA, and dimethyl formamide (DMF). The reaction vessel was purged with nitrogen and then the reaction was allowed to proceed with constant nitrogen flow at 80°C for 18 h. The product was precipitated in cold acetone (5 × 50 mL/g CD-Tos), washed with cold diethyl ether (3 × 100 mL), and dried. The degree of modification was determined by ^1^H NMR.

### Synthesis of Ad-MeHA and CD-MeHA

Methacrylate modification of the HA backbone was achieved via methacrylate esterification with the primary hydroxyl group of sodium HA at basic pH^48,49^. The degree of methacrylate modification was controlled by the amount of methacrylic anhydride introduced during synthesis and was determined to be 28% of the HA disaccharides via ^1^H NMR (**Figure S1**). Next, the methacrylated HA (MeHA) was reacted with proton exchange resin and titrated with TBA-OH to yield methacrylated hyaluronic acid tert-butyl ammonium salt (MeHA-TBA) as previously described^34^ (**Figure S2**). Next, Ad-modified MeHA (Ad-MeHA) and β-CD-modified MeHA (CD-MeHA) were synthesized^49^ by anhydrous coupling. Ad-MeHA was prepared using 1-adamantane acetic acid via di-tert-butyl bicarbonate (BOC_2_O)/4-dimethylaminopyridine (DMAP) esterification while CD-MeHA was prepared using CD-HDA via (benzotriazol-1-yloxy) tris(dimethylamino) phosphonium hexafluorophosphate (BOP) amidation. Products were dialyzed against DI water and degree of modification was determined using ^1^H NMR. Esterification of MeHA with 1-adamantane acetic acid resulted in 43% adamantane modification (by ^1^H-NMR, **Figure S3**) while coupling β-CD-HDA to MeHA via amidation resulted in 26% of the HA disaccharides modified with β-cyclodextrin (by ^1^H-NMR, **Figure S4**).

### Electrospinning

Ad-MeHA, or CD-MeHA, was dissolved at 2% (w/v) in DI water along with 3.5% (w/v) poly(ethylene oxide) (PEO, 900 kDa) and 0.05% (w/v) Irgacure 2959 (I2959) for 24-48 h prior to electrospinning. The polymer solutions were electrospun using a Spraybase (Kildare, Ireland) collection plate set-up and using the following collection parameters: applied voltage: 9.5-10.5 kV, distance from needle to collector: 16 cm, needle gauge: 20, flow rate: 0.4 mL h^−1^. Hydrogel nanofibers were deposited onto foil covering the collector plate, placed into a container which was purged with nitrogen, and crosslinked with UV light (365 nm) for 15 minutes. Parameters were chosen based on previous work with MeHA^50^ and CD-MeHA^51^.

### Nanofiber imaging and quantification

To measure the diameters of the Ad-MeHA and CD-MeHA fibers, samples were electrospun onto foil. After electrospinning, samples were photocrosslinked and analyzed in both dry and swollen states. Dry fibers were imaged using scanning electron microscopy (SEM, FEI Quanta 650) at a magnification of 10,000x. To visualize swollen fibers, a methacrylated rhodamine dye (MeRho, Polysciences, 2 mM) was incorporated prior to electrospinning. Rhodamine-labeled fibers were hydrated, and broken up via trituration through increasingly smaller needle gauges (16G-30G) before encapsulation in a 2% (w/v) MeHA hydrogel. The hydrated fibers were allowed to swell within the MeHA hydrogels overnight in PBS before being imaged using confocal microscopy (Leica inverted confocal microscope, DMi8). Fiber diameters (*n* = 300 fibers per group) were measured from the resulting SEM and confocal images for dry and swollen fibers, respectively, using ImageJ (NIH).

### Hydrogel formulations

Collected guest Ad-MeHA and host CD-MeHA fibers were hydrated (0.1% w/v in DI water) overnight at 37°C to remove PEO from the electrospinning process^52^, centrifuged, supernatant discarded, and then lyophilized. Once dry, the fibers were again hydrated at 0.1% w/v in DI water, allowed to swell at 37°C for at least 2 hours, and then passed through needles of progressively smaller gauge sizes via trituration. Starting with a 16G needle and progressing up to a 30G needle, the fibers were passed through the needles until the fiber solution could easily pass through the needle. This process separated any adjoined fibers and resulted in a reproducible fiber suspension. Fibers were then centrifuged to remove excess water, supernatant discarded, and then resuspended in PBS to achieve the desired fiber concentration. For mechanical testing, the complementary guest and host hydrogel fibers were gently mixed together directly on the rheology plate to create the mixed fibrous guest-host network. Non-fibrous MeHA and guest-host (Ad-MeHA/CD-MeHA) hydrogel groups were prepared as 3% (w/v) solutions and underwent covalent crosslinking (for mechanical stabilization) via photopolymerization in the presence of UV light (365 nm, 10 mW/cm^2^) and 1 mM lithium acylphosphinate (LAP) photoinitiator for 5 min.

### Rheology

All rheological measurements were performed on an Anton Paar MCR 302 rheometer with the plate temperature set at 37°C. The non-fibrous MeHA and guest-host hydrogels were characterized using a cone-plate geometry (25 mm diameter, 0.5°, 25 μm gap) immediately following photopolymerization while guest-host fibrous hydrogels were characterized using a parallel-plate geometry (25 mm diameter, 25 μm gap). Mechanical properties for the non-fibrous MeHA and guest-host hydrogel groups were tested using oscillatory frequency sweeps (0.1−10 Hz, 0.5% strain). Fibrous guest-host hydrogels mechanical properties were tested using oscillatory frequency sweeps (also 0.1−10 Hz, 0.5% strain) and cyclic deformation tests alternating between 0.5% and 250% strain to assess shear-thinning and self-healing capabilities^42^.

### Cell culture

Human mesenchymal stromal cells (hMSCs, Lonza) were used at passage 7 for all experiments. Culture media contained Gibco minimum essential medium (MEM-α) supplemented with 20 v/v% fetal bovine serum (Gibco) and 1 v/v% penicillin/streptomycin/amphotericin B (1000 U/mL, 1000 μg/mL, and 0.25 μg/ mL final concentrations, respectively, Gibco). Prior to cell encapsulation, all materials were modified with thiolated RGD peptide (GCGYGRGDSPG, Genscript) via Michael-type addition in 0.2 M triethanolamine at pH 8, dialyzed against DI water, and then lyophilized. The final RGD concentration was 1 mM for all hydrogel formulations used for cell culture. After materials were put into solution (non-fibrous MeHA and guest-host hydrogels) or hydrated (guest-host fibers), the materials were sterilized using germicidal UV irradiation for 3 h. Prior to the addition of cells, guest-host fibers were centrifuged briefly, with excess liquid aspirated under sterile conditions. To evaluate cell protection during injection under needle flow, hMSCs were added to the non-fibrous hydrogels or hydrogel fibers such that the final encapsulation density was 1 x 10^6^ cells/mL. Materials with cells were loaded into 1 mL tuberculin syringes and hydrogels were injected through a 16G needle at 12 mL h^−1^ using a syringe pump. The extruded material was used to make hydrogel plugs with identical volumes of 40 μL. All groups were subjected to the 5 min UV polymerization to ensure that differences in cell viability were not due to UV cytotoxicity. For all cell culture experiments, media was replaced every 2-3 days.

### Confocal imaging and analysis

Cell viability was assessed with a LIVE/DEAD assay kit (Invitrogen) by incubating hydrogels at 37°C for 30 min with 2 mM calcein-AM and 4 mM ethidium homodimer-1 in PBS plus glucose (G-PBS, glucose 25 mM) followed by three PBS washes. Cells were imaged using a Leica DMi8 inverted confocal microscope and live/dead cells were quantified from 50 μm thick z-stacks (10 μm slices) using ImageJ’s 3D object counter tool. Cell projected area, cell shape index (CSI), and aspect ratio were determined using ImageJ’s measurement tool. CSI determines the circularity of the cell, where a line and a circle have CSI values of 0 and 1, respectively, and was calculated using the formula 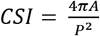 where *A* is the cell projected area and *P* is the cell perimeter. Individual measurements were made for 60 cells from at least 6 hydrogels per experimental group.

### Statistical Analysis

All experimental groups included at least 6 hydrogels for analysis of cell viability and shape. Cell viability and shape data were statistically analyzed using one-way analysis of variance (ANOVA) followed by Tukey’s HSD post-hoc testing. These statistical analyses were conducted using GraphPad Prism 9.0 and R. *P* values < 0.05 were considered statistically significant. For analysis of hydrogel rheological properties, 3 hydrogels were tested per group where the mean values are plotted. Bar graph heights correspond to the mean with standard deviation error bars and individual data points are included as scatter plots overlaying the bars. Tukey box plots of individual cell data show the second and third quartiles as boxes, the median as a line between the boxes, and error bars with the lower value of either 1.5 times the interquartile range or the maximum/minimum value. Data points outside this range are shown individually.

## Results and Discussion

### Guest-host hydrogel fiber design, synthesis, and quantification of fiber diameter

In this study, we report the design, synthesis and characterization of a self-assembling guest-host fibrous hydrogel that can serve as a cell carrier for injectable tissue engineering applications. Hydrophobic Ad (guest) groups associate strongly with the hydrophobic core of CD (host) resulting in a stable supramolecular, yet reversible, guest-host interaction (association constant, K_a_ = 1 x 10^5^ M^−1^)^53^ (**Figure 1**). Ad and CD moieties were separately coupled to methacrylate-modified HA (MeHA)^54^, forming Ad-MeHA and CD-MeHA^51^ respectively (Figure 1A). Modification of the HA backbone with methacrylate groups enables covalent photocrosslinking of the electrospun hydrogel nanofibers in the presence of photoinitiator via UV light-mediated radical polymerization. Following photocrosslinking, the stabilized polymeric fibers will imbibe water upon hydration rather than dissolving^51,55^, creating a water-swollen hydrogel fiber structure. When the hydrated guest and host fibers are mixed, we hypothesized that the complementary nanofibers would associate via hydrophobic supramolecular interactions to form a mechanically robust 3D fibrous hydrogel (Figure 1D) capable of shear-thinning and self-healing. The supramolecularly-assembled fibers can create a hierarchical assembly that provide physical cues to cells at different length scales, mimicking the 3D cues provided by the native fibrous ECM. Additionally, the hydrogel design enables the facile addition of ligands, such as cell adhesion peptides, creating the potential to capture and independently modulate multiple features of native ECM in addition to biophysical cues.

**Figure 1.**
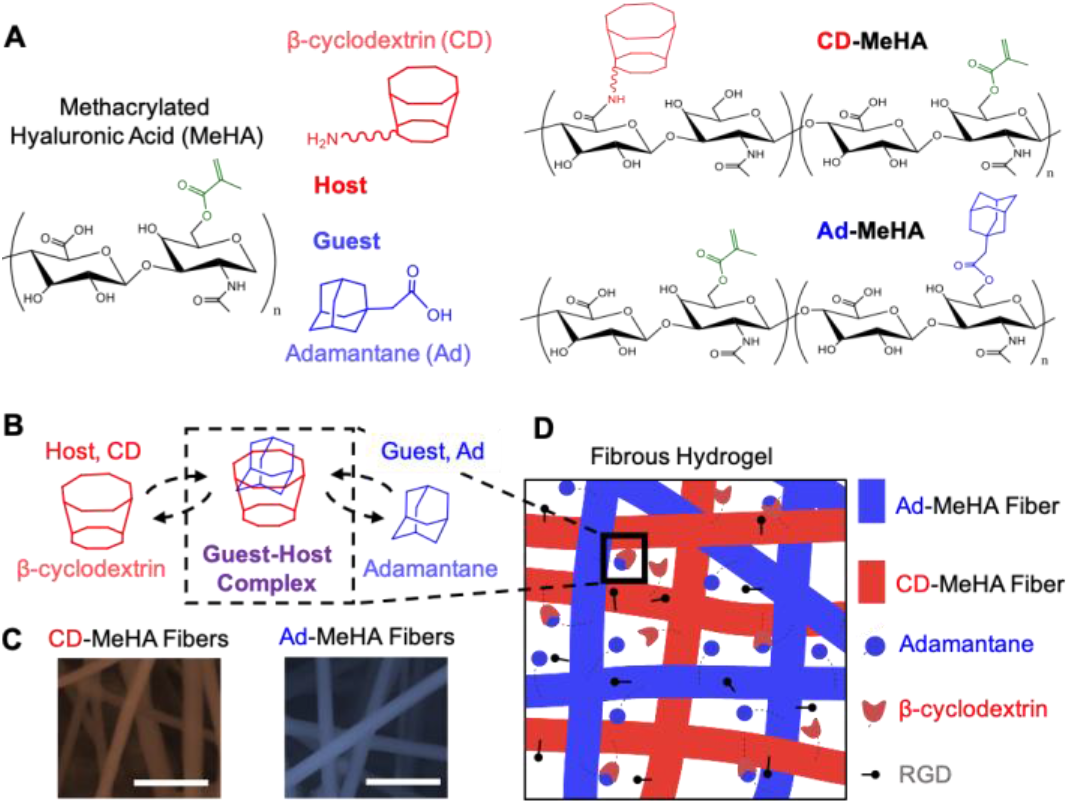
Guest-host supramolecular design to make injectable fibrous hydrogels. A) Structures of guest (Ad-MeHA) and host (CD-MeHA) macromers. The methacrylates enable photocrosslinking to stabilize the fiber structure while B) interaction of β-cyclodextrin (CD, host) and adamantane (Ad, guest) groups forms a reversible guest-host inclusion complex to enable shear-thinning and self-healing. C) SEM of host (CD-MeHA) and guest (Ad-MeHA) electrospun fibers prior to hydration and hydrogel formation. D) Schematic of the fibrous hydrogel composed of mixed guest and host fibers decorated with RGD to permit integrin-mediated cell adhesion. The interactions between Ad and CD on complementary fibers results in assembly of a macroscale fibrous scaffold.

To create the electrospun hydrogel nanofibers capable of supramolecular assembly, Ad-MeHA and CD-MeHA (2% w/v) aqueous polymer solutions were mixed separately with poly(ethylene oxide) (PEO) and the photoinitiator Irgacure 2959 (I2959), and then electrospun to produce guest and host fiber populations (Figure 1C). Addition of the photoinitiator allowed for subsequent stabilization of the fibers by UV light-mediated radical crosslinking of methacrylates while PEO was included as a bioinert carrier polymer. As a carrier polymer, PEO aids in the electrospinning process by making the solution more viscous, thereby disrupting the relatively high surface tension and inducing chain entanglements of the low molecular weight HA solution^52^. Crosslinked hydrogel nanofibers were examined in their dry state via scanning electron microscopy (SEM, **Figure 2A**) and in their hydrated form via confocal microscopy (Figure 2B). Dry Ad-MeHA nanofibers had an average diameter of 234 ± 64 nm while the average dry CD-MeHA fiber diameter was 171 ± 64 nm. Upon hydration, the hydrophilic nanofibers imbibe water, resulting in significant fiber swelling and increased diameter. The hydrated Ad-MeHA nanofibers swelled to an average diameter of 2.16 ± 0.92 μm and hydrated CD-MeHA nanofibers swelled to an average diameter of 1.65 ± 0.54 μm. SEM analysis of native fibrous ECMs has reported fibril diameters in the range of 75-400 nm^56,57^ with small type I collagen fibers in the 1-5 μm range^58^. Therefore, these CD-MeHA and Ad-MeHA hydrogel nanofibers are within the physiologically relevant range for fibrous ECM components^15^. In addition, both the distribution of hydrogel nanofiber diameter^52,59^ and the morphological change of the nanofibers following hydration (and the resulting PEO extraction)^52,60^ have been previously observed in electrospun materials. Likewise, the swelling of the HA-based fibers was anticipated as this same phenomenon was previously reported in electrospun HA matrices^61,62^.

**Figure 2.**
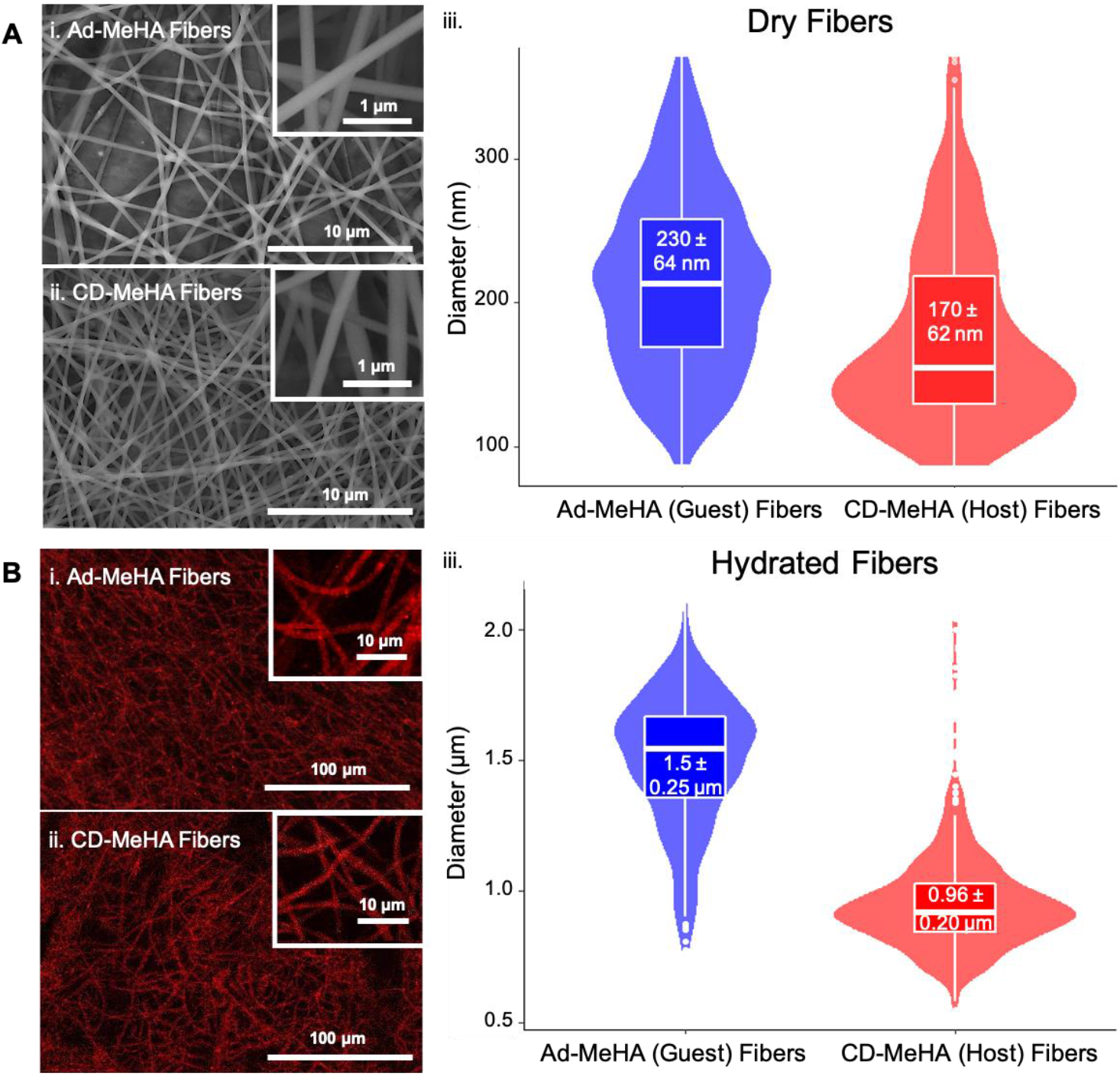
Guest and host fiber morphology and diameter distribution. A. SEM images of photocrosslinked guest (A.i.) and host (A.ii.) fibers prior to hydration with quantification (A.iii.) of dry fiber diameter (*n* = 300 fibers). B. Confocal images of hydrated guest (B.i.) and host (B.ii.) fibers and quantification (B.iii) of the swollen hydrogel fiber diameter (*n* = 300 fibers).

### Guest-host-assembled fibers show mechanical integrity as well as shear-thinning and self-healing character

With the stable structure of the hydrated electrospun hydrogel nanofibers confirmed via confocal microscopy, mechanical properties of the fibrous hydrogels were then characterized via oscillatory shear rheology (**Figure 3**). A 2:1 molar ratio of guest Ad to host CD groups within the fiber mixture was chosen based on previous work with non-fibrous guest-host hydrogels showing that excess Ad groups promoted longer-term mechanical stability^42^. The fiber density of each group in Figure 3 was 1% w/v such that any change in material properties could be directly attributed to guest-host guest interactions. Guest and host fiber populations, measured separately, showed a higher storage modulus than loss modulus as these bulk measurements reflect the properties of the photocrosslinked fibers and their ability to entangle. The guest-host fibrous network also demonstrated a higher storage modulus (G′, 6.6 ± 2.0 kPa) than loss modulus (G″, 1.2 ± 0.5 kPa), but the increase in storage modulus of the mixed guest-host fibers compared to the individual fiber populations highlights the combined mechanical contributions of the individual covalently-crosslinked fibers and the supramolecular interactions between complementary fiber types (Figure 3A).

**Figure 3.**
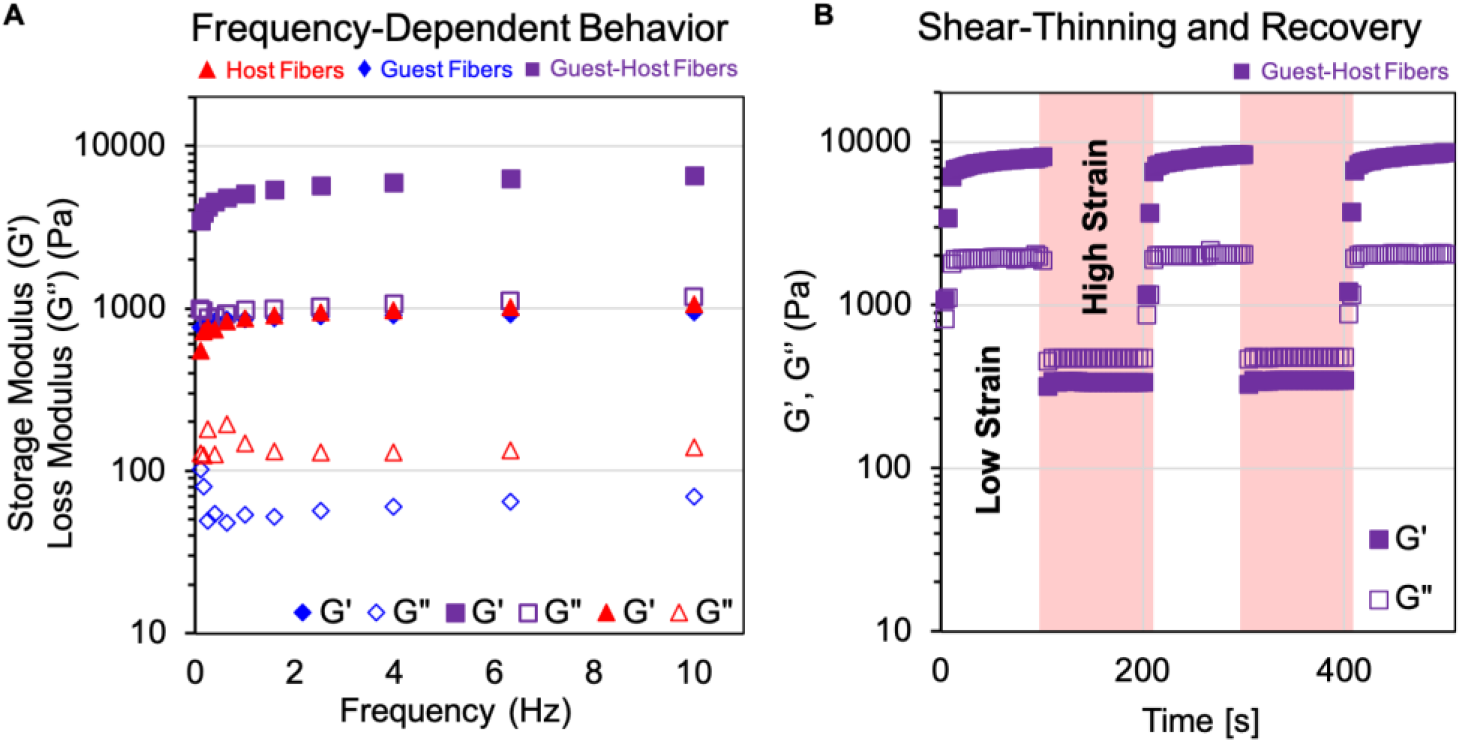
Guest-host-assembled fibers show mechanical integrity as well as shear-thinning and self-healing character. A. Frequency sweep of individual, Ad-MeHA (guest) and CD-MeHA (host), hydrogel fibers and guest-host mixed fibers at constant strain of 0.5%. The storage moduli (G′) of all fiber populations are greater than the loss modulus (G″), reflecting the properties of the photocrosslinked fibers and their ability to entangle. However, the higher moduli of the guest-host hydrogel fiber mixture demonstrates the combined contributions of fiber photocrosslinking and supramolecular interactions between the complementary fiber populations. B. Five-step cyclic deformation test of low strain (0.5%, 100 s) and high strain (250%, 100 s). Guest-host fibrous hydrogels show higher loss moduli than storage moduli at high strains and quickly recover viscoelastic properties at low strains, highlighting shear-thinning and self-healing properties.

Biopolymer density can also be adjusted to tune mechanical properties. Fibrous hydrogel matrices formed with 5% fiber density (also 2:1 Ad:CD molar ratio) showed improved storage moduli over the 1% fiber density formulation (**Figure S5**). These results are consistent with previous work demonstrating that the mechanical properties of guest-host materials are dependent on the quantity of guest species compared to host moieties present in the network^42,63^ and biopolymer density^42^. Other work with electrospun CD-MeHA fibers assessed the guest-host Ad-CD association strength by “gluing” together two CD-MeHA fiber layers with Ad-functionalized HA (AdHA)^51^. This study similarly found that the guest-host pair out performed groups without supramolecular assembly in terms of adhesion strength between the fiber layers. The ability to tune the mechanical properties of the fibrous hydrogel scaffold, by controlling the density of fibers and the ratio of host to guest moieties, is an important feature that may allow access to a range of mechanical properties suitable for fibrous tissue repair.

Having demonstrated the guest-host assembly of the functionalized HA hydrogel fibers, we sought to assess their shear-thinning and self-healing capabilities. The guest-host fibrous scaffold was subjected to cyclic deformations to investigate the responsiveness to shear force and the ability of the fibers to undergo self-healing, i.e., testing the injectable capabilities of the material. Shear-thinning hydrogels will demonstrate sharp decreases in storage and loss moduli when subjected to shear forces with loss moduli crossover (exceeding storage moduli), which is predictive of liquid-like behavior during needle flow upon injection. Conversely, upon cessation of shear, the storage and loss moduli recover for self-healing hydrogels^64^. Multiple cycles of high strain (250%) followed by short recovery periods at low strain (0.5% strain) demonstrated the guest-host fibrous hydrogel’s ability to both shear-thin and self-heal following mechanical deformation (Figure 3B). Based on these experimental findings, we anticipated that the shear forces associated with syringe-based extrusion would induce the same reversible liquid-like to solid-like transitions.

Seminal work using non-fibrous Ad-CD guest-host assembled hydrogels demonstrated the utility of this supramolecular pair, showing tunable mechanics and flow characteristics of the guest-host assembly by altering the guest-host pair ratio and density as well as network structure^42,65,66^. The fibrous hydrogel developed here displays analogous rheological behavior to non-fibrous guest-host hydrogels. As a result of the dynamic bonding interactions between fibers, the guest-host fibrous hydrogel is capable of shear-induced flow (injectability) and rapid recovery. Overall, rheological analysis of the guest-host fibrous hydrogel demonstrated properties of robust mechanical integrity, shear-thinning, and rapid recovery for stability post-injection. Indeed, the fibrous hydrogel scaffold was readily injectable, flowing easily through a needle (12 mL h^−1^, 16G) and recovered as a stable hydrogel plug.

### Injected hMSCs encapsulated in fibrous hydrogels are viable and show increased spreading compared to non-fibrous hydrogels

Finally, we tested the ability of the fibrous guest-host hydrogel to support sustained viability and spreading of encapsulated hMSCs following injection. We chose hMSCs for these experiments due to their multipotential for differentiation toward cell types relevant for a broad range of fibrous tissues^67^ such as muscle, tendon, and ligament^68^. Protecting cells during injection and preserving high viability is one of the fundamental requirements for subsequent therapeutic application, but many injectable delivery vehicles suffer from poor cell survival^69,70^. Previous studies have addressed this issue using materials that leverage physical crosslinking since their compliant mechanical properties support non-uniform network deformation^40,63,71^. In these non-uniform matrices, mechanical stresses imposed on the material do not induce uniform strain fields allowing network deformations to dissipate stress^8^. Therefore, we anticipated that the guest-host fibers could mitigate cell-damaging forces during injection due to the force dissipation capabilities of shear-thinning materials.

We investigated the post-injection viability and spreading of hMSCs when cultured in three hydrogel groups: non-fibrous MeHA, non-fibrous guest-host, and fibrous guest-host (**Figure 4**). Groups were chosen such that the influence of fibrous architecture on cell behavior could be assessed independently from the role of guest-host complexation. All groups contained 1 mM RGD peptide to support cell adhesion and had similar storage moduli (rheology of non-fibrous hydrogels is shown in **Figure S6**). Cell viability was observed at three time points: immediately following injection, day 3 of culture, and day 7 of culture. Following injection, hMSCs in the fibrous guest-host hydrogel were 85 ± 5% viable, a value that was statistically indistinguishable from the hMSCs population encapsulated in either of the non-fibrous hydrogels: MeHA (88 ± 4%) or guest-host (89 ± 3%). Previous studied have reported the highest cell survival in injectable hydrogels (~ 90%) utilizing formulations with modest mechanical properties (G′ ~ 30 Pa)^69,72^. More recently, a self-assembled fibrous peptide hydrogel reported 86.8% cell viability post-injection despite a substantially higher storage modulus (3.1 kPa)^40^. One possible explanation for the fibrous materials’ successful protection of cells during injection, despite the more robust bulk mechanical stiffnesses compared to other successful cell carrier materials, is the stochastic nature of self-assembling fiber hierarchical structures allowing microstructural deformation mechanisms such as shear attenuation via fiber sliding^73–75^. Local shear attenuation is important for native tissue mechanical function and may be mimicked by the supramolecular interactions between complementary guest and host hydrogel fibers, leading to increased force dissipation and thereby protecting encapsulated cells from extensional flow at the entrance of the syringe needle and the subsequent disruption of the cellular membrane^8,69,70^.

**Figure 4.**
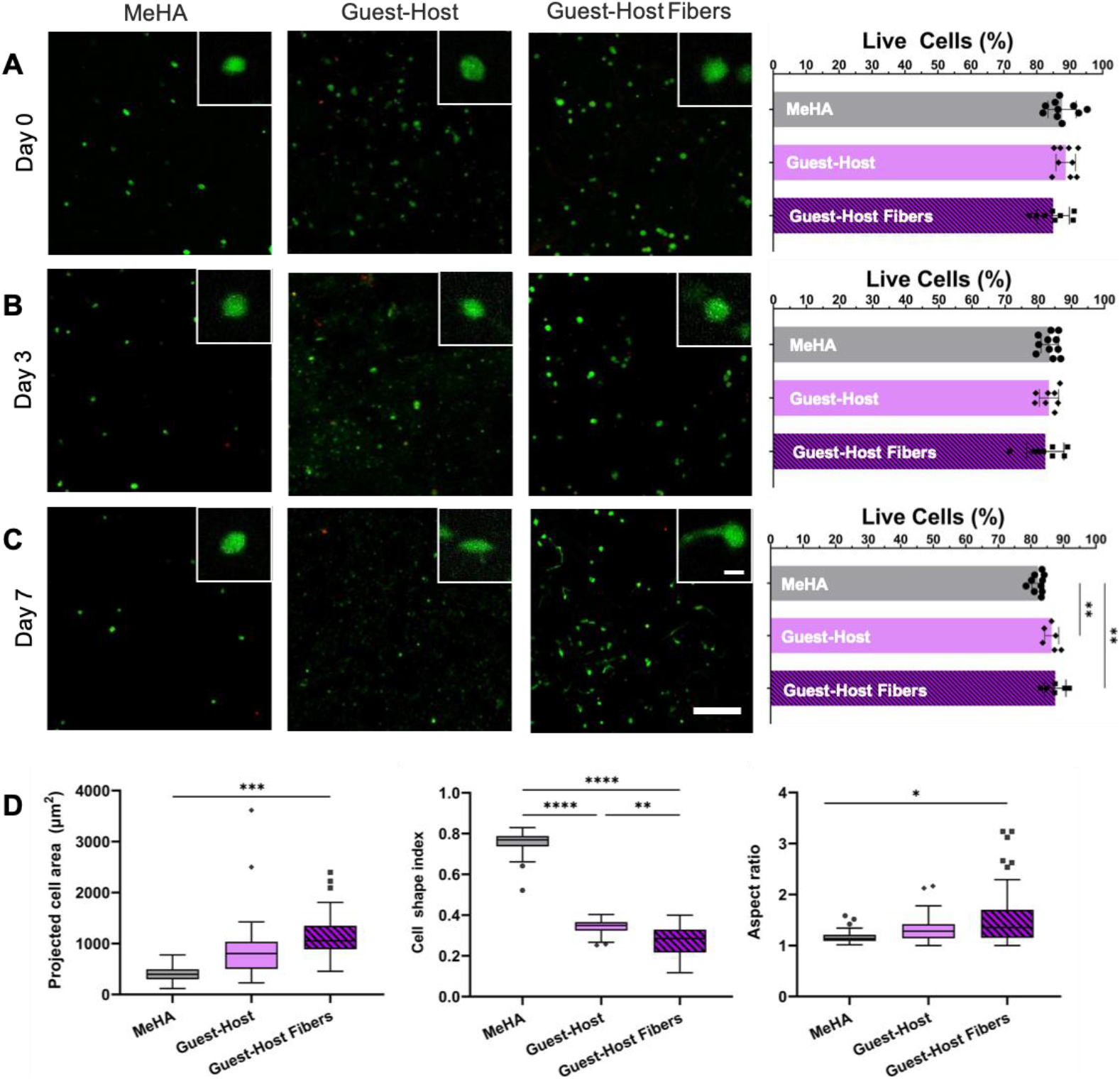
Injected hMSCs encapsulated in fibrous hydrogels are viable and show increased spreading compared to non-fibrous hydrogels. Human mesenchymal stromal cells (hMSCs) were encapsulated in non-fibrous MeHA, non-fibrous guest-host, and fibrous guest-host hydrogels. Live/Dead images of viable (*green*) and membrane damaged (*red*) cells as well as cell viability quantification for all groups is shown A. immediately following injection (Day 0), B. after 3 days of culture, and C. after 7 days of culture. Inset image scale bar = 10 μm. Full image scale bar = 100 μm. Cell viability was comparable across the different hydrogel formulations. D. Quantification of Day 7 cell shape metrics. Cells in fibrous guest-host hydrogels showed significantly increased projected cell area and elongation compared to cells in MeHA hydrogels. Additionally, hMSCs in fibrous guest-host hydrogels showed significantly reduced cell shape index (circularity) compared to hMSCs in non-fibrous guest-host hydrogels. Viability data are presented as mean +/− SD. Tukey box plots of individual cell data (60 cells per group) show the second and third quartiles as boxes, the median as a line between the boxes, and error bars with the lower value of either 1.5 times the interquartile range or the maximum/minimum value. Data points outside this range are shown individually. * *P* < 0.05, ** *P* < 0.01, *** *P*< 0.001, **** *P* < 0.0001. *n* = at least 6 hydrogels per experimental group.

Similar to the initial cell viability, the live cell fractions observed on days 3 and 7 were largely the same (day 3: non-fibrous MeHA: 83 ± 3% and guest-host: 83 ± 3% versus the fibrous guest-host hydrogel: 82 ± 6%), although on day 7 there was a small but statistically significant difference in viability between the non-fibrous MeHA hydrogel group compared to both guest-host hydrogel groups (non-fibrous: 86 ± 2%, fibrous: 87 ± 3%). Although hMSC viability was similar across all experimental groups, qualitative differences in cell shape/spreading observed at day 7 motivated us to quantify hMSC shape metrics such as projected cell area, cell shape index (CSI, a measure of cell circularity), and aspect ratio (Figure 4D). The ability of encapsulated hMSCs to spread is particularly important for supporting the active mechanical signaling^76^ necessary for cells to differentiate^77^ and deposit tissue-specific ECM^78,79^ during fibrous musculoskeletal tissue repair.

hMSCs encapsulated in fibrous guest-host hydrogels showed the highest levels of spreading and elongation (as measured by both CSI and aspect ratio). In particular, the differences in cell shape were the greatest between cells encapsulated in the guest-host hydrogels, both fibrous (spread area: 1130 ± 146 μm^2^, CSI: 0.28 ± 0.04, aspect ratio: 1.54 ± 0.33) and non-fibrous (spread area: 850 ± 333 μm^2^, CSI: 0.34 ± 0.02, aspect ratio: 1.33 ± 0.09) compared to the MeHA hydrogels which showed more rounded hMSC morphologies (spread area: 400 ± 93 μm^2^, CSI: 0.76 ± 0.02, aspect ratio: 1.17 ± 0.05). The reduction in hMSC spreading and elongation found in MeHA hydrogels compared to the guest-host hydrogels is likely due to differences in viscoelasticity. The guest-host hydrogel networks contain both covalent crosslinking and guest-host supramolecular interactions, leading to viscoelastic properties as shown in our rheological analysis while MeHA hydrogels are covalently crosslinked and behave like elastic solids. Numerous recent studies, including one using the Ad-CD guest-host pair^80^, highlighted the role of hydrogel viscoelasticity in supporting increased cell spreading and elongation in 3D cultures^79,81^. Our findings, in particular the significant reduction in CSI observed for hMSCs in fibrous guest-host hydrogels compared to non-fibrous guest host hydrogels, are also supported by recent work demonstrating that human fibroblasts show increased outgrowth and spreading within fiber-containing hydrogel composites compared to non-fibrous hydrogels^28^. Together, the data from these experiments underscore the suitability of our fibrous hydrogel design for injectable encapsulation of hMSCs in a microenvironment supportive of cell spreading and elongation.

## Conclusions

In summary, we designed and fabricated an injectable fibrous hydrogel capable of shear-thinning and self-healing in physiological conditions. Using HA as a naturally-derived polymer already applied in clinical settings for decades, these materials offer unique advantages with respect to biocompatibility and mechanical integrity (G’ = 6.6 ± 2.0 kPa) while also recapitulating the fibrous architecture of tissue. In addition to exhibiting shear-thinning and self-healing behavior, the guest-host fibrous hydrogel demonstrated injectability wherein encapsulated hMSCs were protected from membrane-disrupting shear forces during injection, resulting in sustained 3D hMSC viability (> 85%). Cells were also more spread and elongated in viscoelastic guest-host hydrogels compared to non-fibrous elastic hydrogel controls, with hMSCs showing significantly reduced CSI (circularity) in fibrous guest-host hydrogels compared to both non-fibrous MeHA and non-fibrous guest-host hydrogels. In summary, the injectable fibrous hydrogel platform introduced here offers the potential to broaden minimally invasive delivery to musculoskeletal tissue engineering applications requiring robust structural properties while also laying the foundation for future opportunities in 3D bioprinting and fundamental studies of cell-microenvironment interactions.

## Supporting Information

^1^H NMR spectra for MeHA, Ad-MeHA and CD-MeHA as well as additional hydrogel mechanical characterization can be found in the Supporting Information.

## Acknowledgments

We thank Prof. Rachel Letteri and Dr. Matthew D. Davidson for helpful discussions as well as Erica Hui for providing initial hydrogel synthesis training. Utilization of the FEI Quanta 650 SEM within UVA’s Nanoscale Materials Characterization Facility (NMCF) was fundamental to this work and we thank the staff for equipment training. This work was supported by the University of Virginia, the DoD (W81XWH-19-1-0157), and the NIH (R21AR075181). The content is solely the responsibility of the authors and does not necessarily represent the official views of the National Institutes of Health.

