## Supplemental Figures for "Guest-host supramolecular assembly of injectable hydrogel fibers for cell encapsulation"

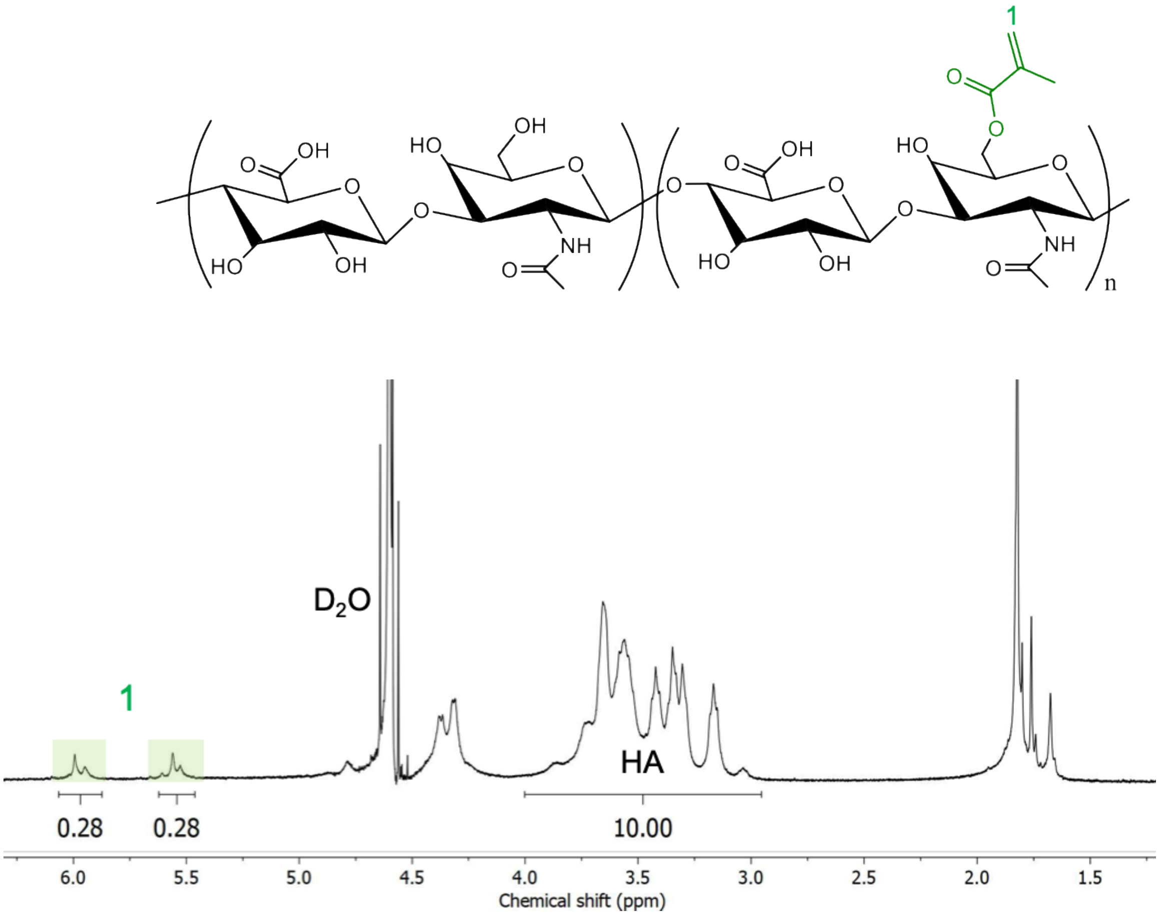


**Figure S1. $^1$H NMR spectrum of methacrylate-modified hyaluronic acid (MeHA)**. The degree of HA modification with methacrylates was determined to be 28%. Degree of functionalization was determined from integration of the methylene (δ = 5.82, 1H and δ = 6.25, 1H) highlighted green and labeled '1' relative to the HA backbone, (δ = 3.10-4.10, 10H).

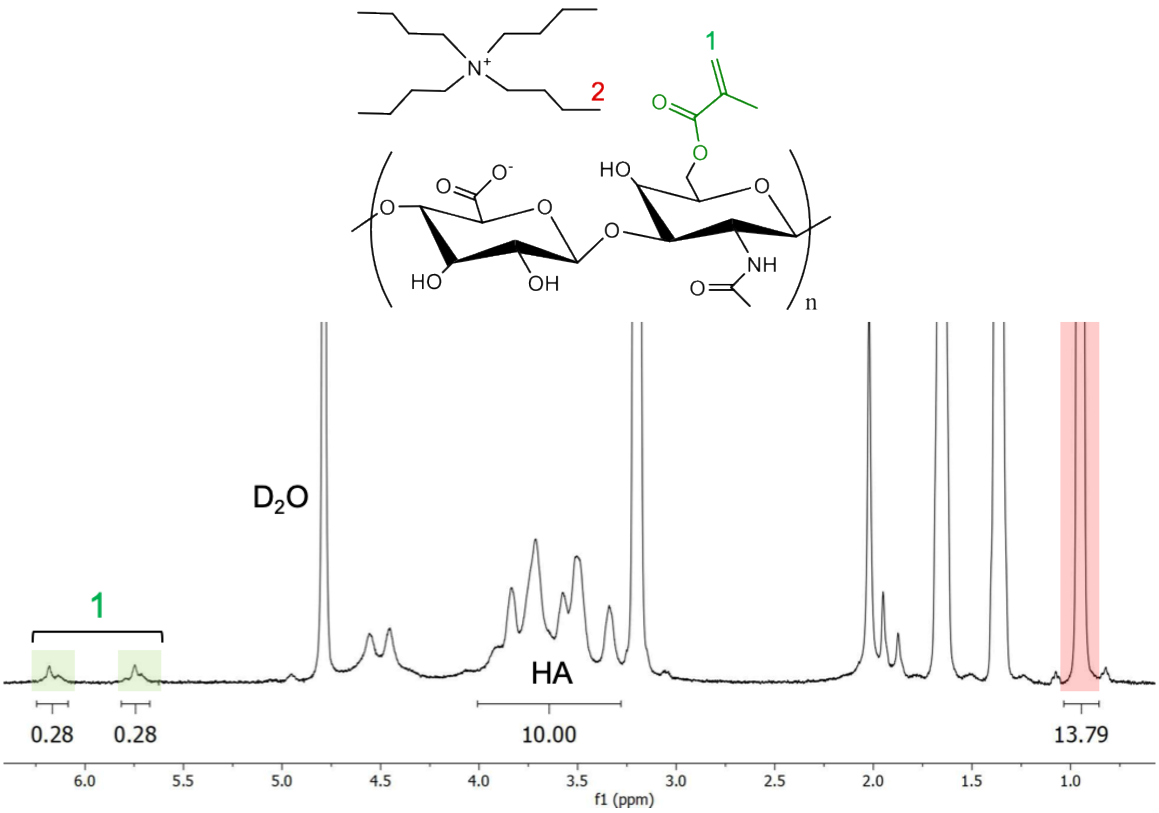


**Figure S2. $^1$H NMR spectrum of methacrylated hyaluronic acid tert-butyl ammonium salt (MeHA-TBA).** The MeHA used for the synthesis of Ad (guest) and CD (host) derivatives underwent addition of the TBA salt (highlighted red) to enable further HA modification.

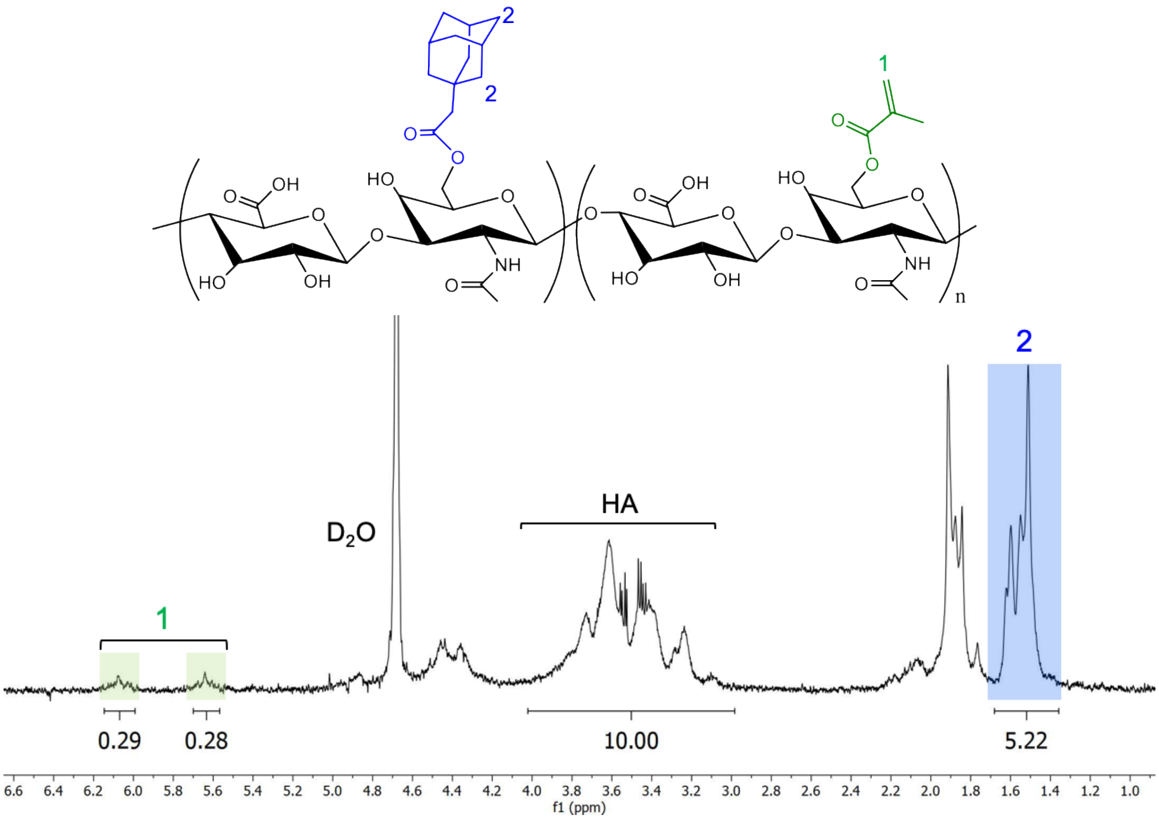


**Figure S3. $^1$H NMR spectrum of adamantane and methacrylate-modified hyaluronic acid (Ad-MeHA).** The degree of MeHA modification with adamantane was determined to be 43%. The integration of the ethyl multiplet of adamantane (δ = 1.40-1.70, 12H) highlighted in blue gives the degree of modification relative to the HA backbone (δ = 3.10-4.10, 10H).

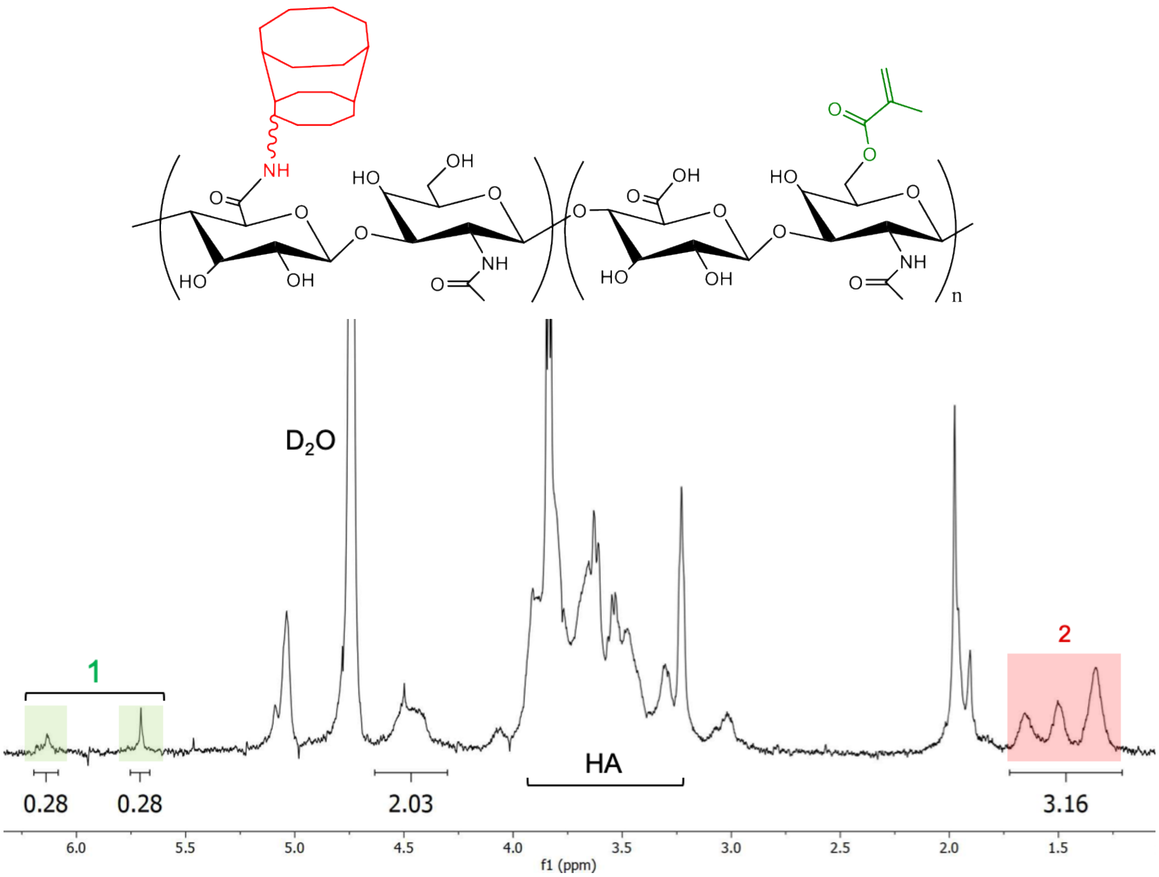


**Figure S4. $^1$H NMR spectrum of β-cyclodextrin and methacrylate-modified hyaluronic acid (CD-MeHA).** The degree of MeHA modification with β-cyclodextrin was determined to be 26%. This was calculated from the integration of the hexane linker (δ = 1.20-1.75, 12H) highlighted in red relative to the methacrylate modification, which is assumed to stay constant.

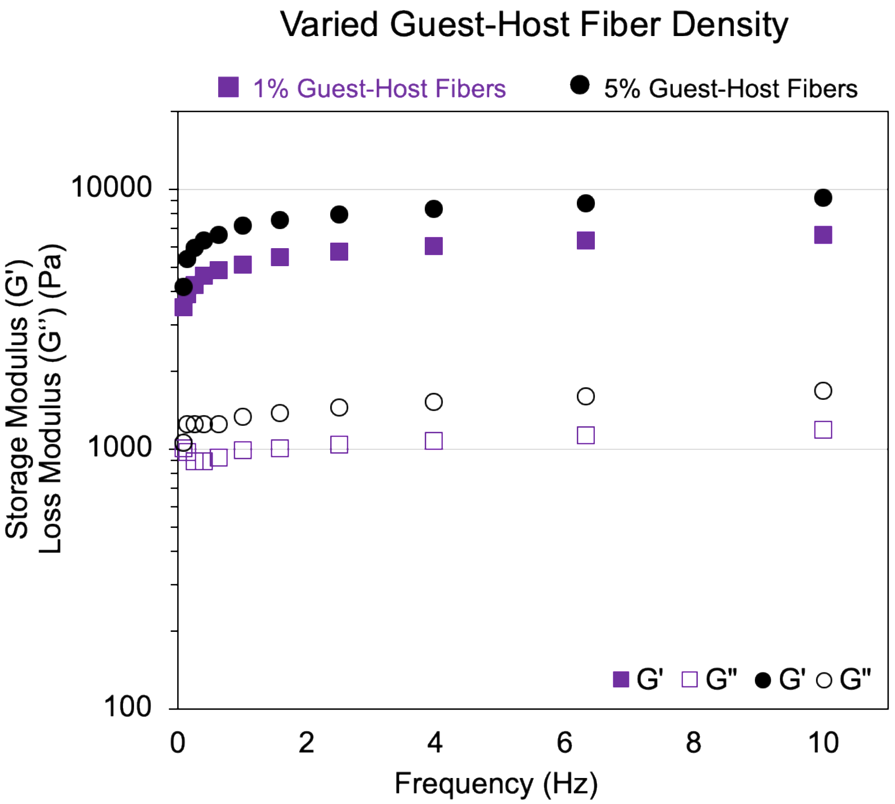


**Figure S5. Rheological properties of the guest-host fiber network with varying fiber density**. The frequency-dependent behavior was measured using a constant strain of 0.5%. The 1% fibrous hydrogel reached a final storage modulus (G′) of 6.6 kPa and the 5% fibrous hydrogel a G′ of 9.2 kPa at 10 Hz.

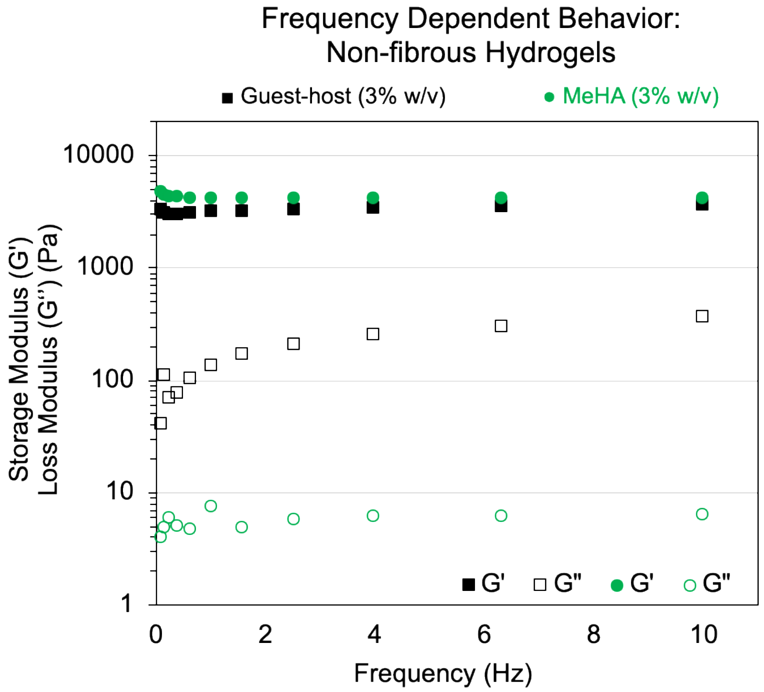


**Figure S6**. **Rheological properties of the non-fibrous MeHA and guest-host hydrogels used for cell encapsulation.** The frequency-dependent behavior was measured using a constant strain of 0.5%. The non-fibrous 3% MeHA hydrogel formulation reached a final storage modulus (G′) of 4.2 kPa and the non-fibrous 3% guest-host hydrogel formulation reached a final G′ of 3.7 kPa at 10 Hz.
